# AFM-based High-Throughput Nanomechanical Screening of Single Extracellular Vesicles

**DOI:** 10.1101/854539

**Authors:** Andrea Ridolfi, Marco Brucale, Costanza Montis, Lucrezia Caselli, Lucia Paolini, Anne Borup, Anders T. Boysen, Francesca Loria, Martijn J. C. van Herwijnen, Marije Kleinjan, Peter Nejsum, Natasa Zarovni, Marca H. M. Wauben, Debora Berti, Paolo Bergese, Francesco Valle

**Affiliations:** Consorzio Interuniversitario per lo Sviluppo dei Sistemi a Grande Interfase (CSGI), Firenze, Italy; Consiglio Nazionale delle Ricerche, Istituto per lo Studio dei Materiali Nanostrutturati (CNR-ISMN), Bologna, Italy; Dipartimento di Chimica “Ugo Schiff”, Università degli Studi di Firenze, Firenze, Italy; Dipartimento di Medicina Molecolare e Traslazionale, Università degli Studi di Brescia, Brescia, Italy; Department of Clinical Medicine, Faculty of Health, Aarhus University, Aarhus, Denmark; HansaBiomed Life Sciences, Tallinn, Estonia; Department of Biochemistry & Cell Biology, Faculty of Veterinary Medicine, Utrecht University, Utrecht, The Netherlands

## Abstract

We herein describe an Atomic Force Microscopy (AFM)-based experimental procedure which allows the simultaneous mechanical and morphological characterization of several hundred individual nanosized vesicles within the hour timescale.

When deposited on a flat rigid surface from aqueous solution, vesicles are deformed by adhesion forces into oblate spheroids whose geometry is a direct consequence of their mechanical stiffness. AFM image analysis can be used to quantitatively measure the contact angle of individual vesicles, which is a size-independent descriptor of their deformation and, consequently, of their stiffness. The same geometrical measurements can be used to infer vesicle diameter in its original, spherical shape.

We demonstrate the applicability of the proposed approach to natural vesicles obtained from different sources, recovering their size and stiffness distributions by simple AFM imaging in liquid. We show how the combined EV stiffness/size readout is able to discriminate between subpopulations of vesicular and non-vesicular objects in the same sample, and between populations of vesicles with similar sizes but different mechanical characteristics. We also discuss a force spectroscopy calibration procedure to quantitatively link the stiffness of EVs to their average contact angle.

Finally, we discuss expected extensions and applications of the methodology.

## Introduction

Extracellular vesicles (EVs) are cell-released, sub-micron membranous particles involved in numerous physiological and pathological functions [van Niel 2018; Yáñez-Mó 2015]. Due to their almost ubiquitous relevance, they are focalizing the interest of a rapidly-growing, highly multidisciplinary research community including oncologists, neurologists, bioengineers, parasitologists, cell biologists, food scientists, and biophysicists [Xu 2018; Vescovi 2019; Galieva 2019; Ostenfeld 2014; Mardahl 2019; van Herwijnen 2018; Roy 2018]. Because of the diverse biogenesis/release mechanisms of EVs and their enormous heterogeneity, the EV-community is making a continuous effort to reach a consensus regarding several fundamental issues, including EV nomenclature [Thery 2018].

The vast majority of experimental research on EVs of any type starts with their isolation, purification and enrichment-which are non-trivial endeavors, often needing sample-specific protocol optimization to limit contamination by non-vesicular material or excessive EV size polydispersion [Jeppesen 2014; Cocucci 2015; Montis 2017; Shao 2018]. Furthermore, the analysis of EV samples is made difficult by a general scarcity of established tools for characterizing EVs with highly varied size, origin, function, membrane lipid/protein composition, and cargo content [Paolini 2018; Thery 2018]. Hence, there is a need to develop methods for rapid, label-free assessment of highly diverse EV-containing samples, able to discern between vesicular and non-vesicular particles in the submicron range. In this context, single-vesicle measurements seem especially promising [Chiang 2019].

One relatively constant feature of EVs isolated from different sources is their mechanical behavior, which is known to be influential to cellular adhesion, endo/exocytosis, cellular uptake and mechanosensing [Sorkin 2018]. EVs have been shown to give a characteristic mechanical response to an applied load: a highly linear force/distance elastic deformation regime, which is also typical of synthetic liposomes but is otherwise very uncommon in non-vesicular objects [Sharma 2010; Calò 2014; Parisse 2017; Vorselen 2018]. This characteristic behavior can be recognized by probing the mechanical response of individual vesicles deposited on a substrate via Atomic Force Microscope (AFM)-based Force Spectroscopy (FS) [Krieg 2019]. The linear deformation regime slope reflects the vesicle’s overall stiffness (k_S_), *i.e*. its resistance to deformation, and can be quantitatively measured via AFM-FS nanoindentation experiments. Specific types of EV were observed to have characteristic k_S_ values, which can vary in presence of pathological processes [Vorselen 2018; Whitehead 2015]. Due to this, it seems reasonable to consider mechanical response in general, and k_S_ in particular, as the basis for a method capable of discriminating EVs from contaminants, or even between different types of EVs.

The observed linear mechanical response of vesicles is best rationalized by the Canham-Helfrich (CH) model [Canham 1970; Helfrich 1973], in which the overall stiffness k_S_ is the sum of two contributing factors: membrane rigidity, quantified by its bending modulus (κ), and luminal pressurization (Π). Wuite and coworkers recently demonstrated that AFM-FS can be employed to separately determine the κ and Π values of individual liposomes [Vorselen 2017], that the same approach is applicable to EVs [Vorselen 2018], and that it can detect quantitative mechanical behavior variations linked to biological function [Sorkin 2018]. This elegant and powerful AFM-FS approach is however quite labor-intensive, requiring the experimental determination of k_S_, tether elongation force (F_T_) and curvature radius (R_C_) for each individual vesicle. In particular, obtaining clear F_T_ readings involves the establishment of a single mechanical link between the vesicle’s membrane and the AFM probe, and can be problematic on EVs with abundant membrane proteins and/or lipopolysaccharides content. Finally, it is necessary to pool the readings of at least several tens of individual vesicles to obtain a reasonably clear picture of an EVs population’s overall mechanical characteristics. Combined together, these considerations imply that the FS-based strategy mentioned above is in our opinion the best currently available method to obtain a quantitative mechanical characterization of individual vesicles, but is also poorly suited to a quick, routine screening of unknown EV samples mainly aimed at achieving a broad picture of their size distribution and purity.

We herein propose a method for the rapid nanomechanical assessment of EV populations based on simple AFM imaging performed in liquid and successive morphometric analysis easily performed with freely available software. Following the procedure detailed in the following sections, it is possible to define the size and mechanical characteristics of a few hundred individual vesicles in the hour timescale in ideal experimental conditions. Although the mechanical readout provided by our procedure is semi-quantitative, it is able to discriminate between subpopulations of vesicular and non-vesicular objects deposited on the same substrate, and between populations of vesicles with similar sizes but different mechanical characteristics. Moreover, we show a calibration procedure that can be used to estimate the k_S_ of EVs without performing FS experiments.

## Materials and Methods

### Liposomes Preparation and Characterization

Different lipids with PC polar headgroup (DOPC (1,2-dioleoyl-sn-glycero-3-phosphocholine), POPC (1-palmitoyl-2-oleoyl-glycero-3-phosphocholine), DPPC (1,2-dipalmitoyl-sn-glycero-3-phosphocholine), DSPC (1,2-1,2-distearoyl-sn-glycero-3-phosphocholine)) were purchased from Sigma Aldrich (St. Louis, MO, USA); lipid dry powders were dispersed in defined amounts of chloroform, to prepare stock solutions. Lipid films were obtained by evaporating appropriate amounts of lipid stock solutions in chloroform under a stream of nitrogen, followed by overnight drying under vacuum. The films were swollen by suspension in warm (50 °C) milliQ water to a final lipid concentration of 4 mg/mL, followed by vigorous vortex mixing. The dispersions were then tip-sonicated for 15min to obtain a dispersion of unilamellar lipid vesicles with low size polydispersity. The size distribution and Zeta Potential of the vesicles was determined through Dynamic Light Scattering and Zeta Potential measurements, respectively (see Figure S4).

### Natural Vesicles Isolation and Purification

All EV data were acquired and reported following MISEV 2018 [Thery 2018] and MIRABEL [Faria 2018] international guidelines. Relevant data were also submitted to the EV-TRACK knowledge base (EV-TRACK ID: EV190077) [Van Deun 2017].

### EVs from bovine milk

Raw milk (100 ml) was collected from the cooled tank from a local dairy farm (Tolakker, Utrecht, The Netherlands), transferred to 50 ml polypropylene tubes and centrifuged for 10 minutes at 22°C at 3000 xg (Beckman Coulter Allegra X-12R, Fullerton, CA, USA). After removal of the cream layer, the milk supernatant was harvested without disturbing the pellet and transferred to new tubes. A second centrifugation step at 3000 xg followed, after which the milk supernatant was collected and stored at −80°C until further processing. Thawed milk supernatant (80 ml) was transferred to polyallomer SW40 tubes (Beckman Coulter) and centrifuged at 5000 xg for 30 minutes at 4°C and subsequently at 10000 xg (Beckman Coulter Optima L-90K with a SW40Ti rotor). For the precipitation of caseins, the milk supernatant was acidified to pH4.6 by adding Hydrochloric acid (HCL, 1M) while stirring. Caseins were pelleted by centrifugation at 360 xg (Beckman Coulter Allegra X-12R) after which casein-free milk supernatant was collected. Next, 6.5 ml of the milk 10000 xg supernatant was loaded on top of a 60% – 10% Optiprep gradient (Optiprep™, Progen Biotechnik GmbH, Heidelberg, Germany) made in a SW40 tube. Gradients were ultracentrifuged at 197000 xg (Beckman Coulter Optima L-90K with a SW40Ti rotor) for 15-18 hrs. After centrifugation, fractions of 500 μl were harvested and densities were measured in order to identify the EV-containing fractions with 1.06-1.19 g/ml, which were pooled. Optiprep was exchanged for PBS by using size exclusion chromatography on the EV-containing fractions pooled in a 20 ml column (Bio-Rad Laboratories, Hercules, CA, USA) packed with 15 ml Sephadex g100 (Sigma-Aldrich, St. Louis, MO, USA). Fractions of 1 ml with eluted from the column by washing with PBS (Gibco™, Invitrogen, Carlsbad, CA, USA). Eluates 3 to 9 were pooled as these contained EVs and samples were stored at −80°C until use.

### EVs from Ascaris suum

Live adult *Ascaris suum* nematodes were obtained from pigs slaughtered at the Danish Crown abattoir in Herning, Denmark. Five worms, two males and three females, were put in a T175 flask and washed in 175 ml RPMI-1640 with 1X Antibiotic-Antimycotic (Thermo Fisher Scientific, cat#15240062) and 1 μg/ml ciprofloxacin (Sigma, cat#17850) (*RPMI-Anti/Anti*) in a total of four cycles of 15 minutes followed by three cycles of one hour of incubation at 37 °C. After washing, the worms were incubated in 175 ml RPMI-Anti/Anti for 72h in a 5% CO_2_ incubator at 37 °C. The media containing excretory/secretory (ES) products from the worms was exchanged and collected every 24 hours. The collected ES products were stored at −80 °C. ES products from all three days were thawed at 4 °C and pooled to be concentrated 720 times with Amicon^®^ Ultra-15 Centrifugal Filter Unit 10 kDa cut-off (Merck, cat#UFC901024). The concentrate was used for EV separation.

To separate EVs, two different methods were used: ultracentrifugation (UC) and size exclusion chromatography (SEC). Ultracentrifugation procedure: 500 μl of the concentrated ES products were transferred to polycarbonate ultracentrifuge tubes with cap assembly (Beckman Coulter, cat#355603) and diluted with PBS 1X to a final volume of 10 ml. Total volume was centrifuged at 10000 xg for 30 minutes at 4°C at 10000 xg for 30 minutes at 4 °C (Beckman Coulter Optima L-80 XP Ultracentrifuge, TI 50 rotor kept at 4°). Supernatant (approx. 10 ml) was transferred to a new polycarbonate ultracentrifuge tube and centrifuged at 100000 xg for 70 minutes at 4°C (Beckman Coulter Optima L-80 XP Ultracentrifuge, TI 50 rotor kept at 4°C). The pellet was then dissolved in 10 ml of PBS 1X and re-centrifuged at 100000 xg for 70 minutes at 4°C. Final pellet was resuspended in 2 ml PBS 1X, transferred to an Eppendorf tube and stored at −80°C. SEC procedure: EVs were separated using qEVoriginal/70 nm columns from iZON (iZON Science Ltd, cat#SP1) according to manufacturing instructions using PBS 1X as buffer. Twenty-four fractions of 500 μl were collected. The fractions 7-10 were pooled as EV-containing fraction and stored at −80°C.

### EV characterization

EV preparations from bovine milk and *Ascaris suum* were characterized for purity from protein contaminants and titrated by Colorimetric Nanoplasmonic Assay (CONAN) assay (Supplementary Table ST1). EV size distribution was in addition determined by Nanoparticle Tracking Analysis (NTA) for samples from Ascaris suum E/S (Supplementary Figures S5-S7). Protein composition was analyzed by Western blot. It is to note that the biochemical characterization can be performed only on bovine milk derived EVs, since no specific protein markers have been identified for *Ascaris suum* samples so far. The presence of EV-associated markers, and non-EV markers is presented in Supplementary Figure S8. Characterization protocol details and results are presented in the Supplementary information file.

### Surface Preparation and Sample Deposition

All AFM experiments were performed on poly-L-lysine (PLL) coated glass coverslips. All reagents were acquired from Sigma-Aldrich Inc (www.sigmaaldrich.com) unless otherwise stated. Microscopy glass slides (15mm diameter round coverslips, Menzel Gläser) were cleaned in a sonicator bath (Elmasonic Elma S30H) for 30 minutes in acetone, followed by 30 minutes in isopropanol and 30 minutes in ultrapure water (Millipore Simplicity UV). Clean slides were incubated overnight in a 0.0001% (w/v) PLL solution at room temperature, thoroughly rinsed with ultrapure water and dried with nitrogen. The water contact angle (1μl droplets at ~25°C, measured with a GBX DigiDrop goniometer) of slides was 26°±1° prior to functionalization and 20°±2° after PLL deposition.

A 10 μl-droplet of the vesicle-containing solution under study was deposited on a PLL-functionalized glass slide and left to adsorb for 10 minutes at 4°C, then inserted in the AFM fluid cell (see below) without further rinsing. The concentration of each vesicle-containing solution was adjusted by trial and error in successive depositions in order to maximize the surface density of isolated, individual vesicles and minimize clusters of adjoining vesicles.

### AFM setup

All AFM experiments were performed in ultrapure water at room temperature on a Bruker Multimode8 (equipped with Nanoscope V electronics, a sealed fluid cell and a type JV piezoelectric scanner) using Bruker SNL-A probes (triangular cantilever, nominal tip curvature radius 2-12 nm, nominal elastic constant 0.35 N/m) calibrated with the thermal noise method [Hutter 1993].

### AFM Imaging

Imaging was performed in PeakForce mode. In order to minimize vesicle deformation or rupture upon interaction with the probe, the applied force setpoint was kept in the 150-250 pN range. Lateral probe velocity was not allowed to exceed 5μm/s. Feedback gain was set at higher values than those usually employed for optimal image quality in order to ensure minimal probe-induced vesicle deformation upon lateral contact along the fast scan axis.

This type of parameter optimization resulted in images with comparatively high noise levels in the empty areas of the surface (≤20nm peak to peak), but in which the height profiles of individual vesicles measured along both the slow and the fast scan axis could be fitted extremely well with circular arcs (Figure S1c). The average height value of all bare substrate zones was taken as the baseline zero height reference.

Image background subtraction was performed using Gwyddion 2.53 [Nečas 2012]. Image analysis was performed with a combination of Gwyddion and custom Python scripts, but it can be easily carried out manually by only using functions included in Gwyddion and a spreadsheet.

### AFM force spectroscopy

The mechanical characterization of vesicles via AFM force spectroscopy was performed following the approach recently described by Vorselen et al [Vorselen 2017]. The sample was first scanned (see previous paragraph) to locate individual vesicles (Figure S1a). The chosen vesicle was then imaged (Figure S1b) at higher resolution (~500×500 nm scan, 512×512 points); its height profile along the slow scan axis was fitted with a circular arc only taking into account values 10nm above the bare substrate (typical fit R^2^ ≥ 0.95). This procedure yielded, for each vesicle, an apparent fitted curvature radius R_C_ and a vesicle height value H_S_ (Figure S1c), which were corrected as described in [Vorselen 2017].

In principle, it would be sufficient to record the force/distance plot of just one approach/retraction cycle for each vesicle measured at its highest point, while avoiding membrane puncturing. In our hands however, this was practically impossible due to intrinsic piezo inaccuracy and drift, which imply a certain degree of uncertainty on both the XY position at which the force curve is performed relative to the original image, and on the maximum applied force.

To overcome this limitation, we recorded a series of force/distance curves at multiple XY positions (typically around 64-100 curves arranged in a square array covering the vesicle initial location Figure S1b, green crosses) for each individual vesicle. In most cases, only a few curves showed the full mechanical fingerprint of an intact vesicle on both the approach and retraction cycles (Figure S1d), showing a linear deformation upon applied pressure and a tether elongation plateau during probe retraction. Of these, we first discarded those with probe-vesicle contact points (P_C_) occurring at probe-surface distances below vesicle height as measured by imaging (P_C_ < H_S_, see previous paragraph). We then discarded traces in which the tether elongation plateau occurring during probe retraction did not extend beyond initial contact point. However, we relaxed this requirement for those natural vesicle samples on which obtaining clean tether plateaus was nearly impossible (see results and discussion section).

Remaining traces (typically 1-3 per vesicle) were analyzed to calculate vesicle stiffness (k_S_) and tether elongation force (F_T_). Multiple valid curves referring to the same vesicle resulted in very narrow distributions of both k_S_ and F_T_ (with average measured values taken as representative for each vesicle), while different vesicles of the same type showed much larger variations (see below). Membrane bending modulus (κ) and internal pressurization (Π) values were then calculated for each individual vesicle using its R_C_, k_S_ and F_T_ values as described in [Vorselen 2017].

## Results and Discussion

### Quantitative AFM morphometry of vesicles

The mechanical characterization of vesicles via quantitative AFM morphometry was performed as follows. Representative AFM micrographs (typically 5×5 μm, 512×512 points) were first acquired as described above. Since all the following image analysis steps rely on a correct zero-height baseline assignment, special care was taken to ensure that the image was devoid of image flattening artifacts by masking all positive features appearing on the surface and excluding them from linear background interpolation. In some cases, it was necessary to iterate the masking/subtraction procedure several times to obtain the required background flatness.

Figure 1a exemplifies a correctly processed AFM image of DPPC liposomes: after background subtraction, height profiles measured along the diagonals of the whole image (Figure 1b) are extremely flat and the average height of empty areas is zero. Moreover, height profiles measured along the X and Y axis for individual vesicles are symmetrical and almost superimposable (Figure 1c), denoting that probe-induced deformation of vesicles along the fast scan axis is marginal.

**Figure 1.**
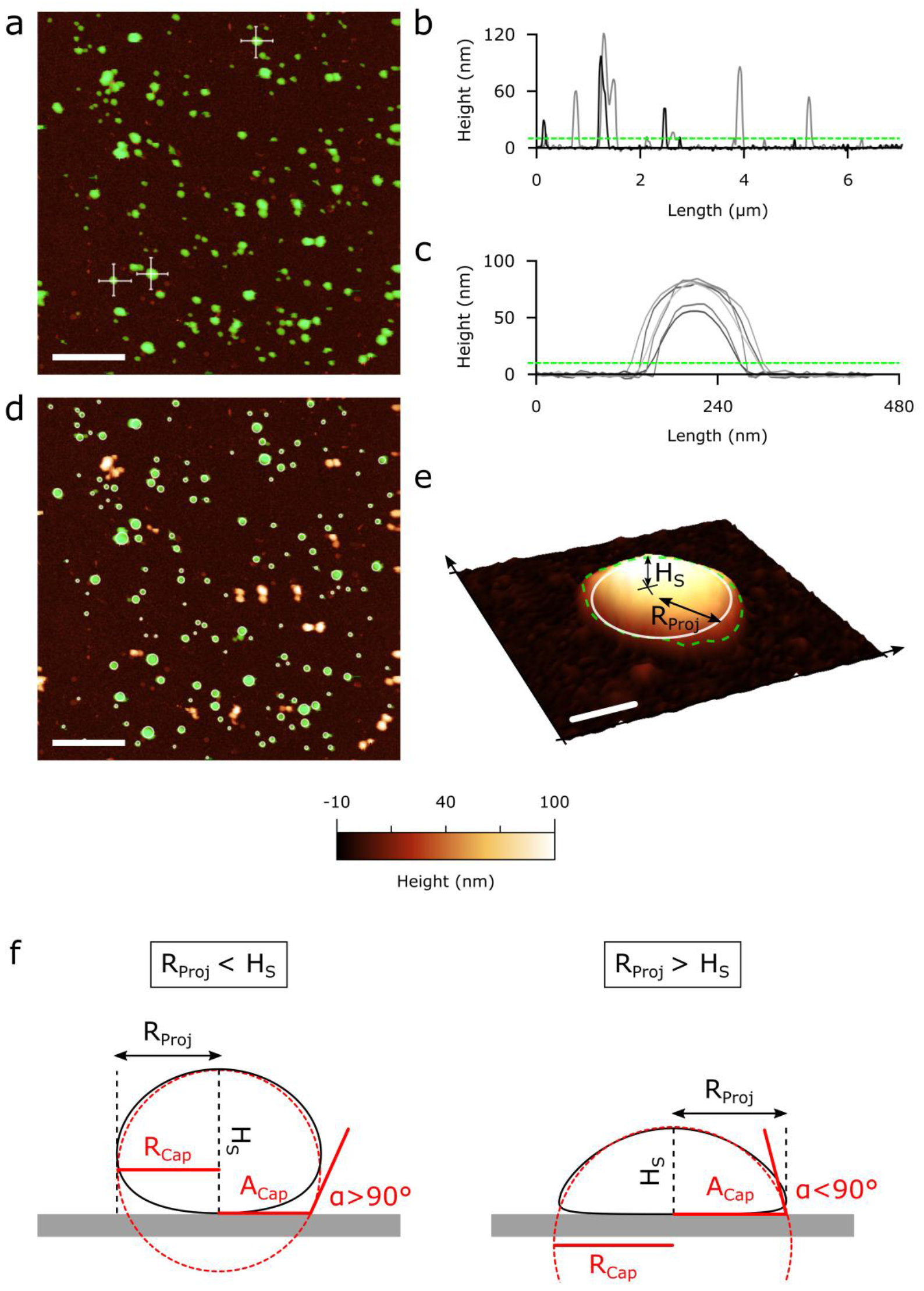
AFM imaging and morphometry analysis. (a): representative AFM micrograph of a DPPC liposome sample deposited on a PLL-functionalized substrate. Scalebar = 1μm. Correct background subtraction is crucial to successive image analysis steps (see materials and methods) and is first checked by plotting height profiles measured along the two diagonals of the whole image (b): after proper flattening, diagonal profiles must show a flat baseline centered at height=0 with positive features. To minimize probe-induced vesicle deformation, imaging should be preferentially performed at low applied load (<250 pN) and high feedback gain (see materials and methods). (c): Profiles of putative vesicles measured along fast and slow scan axes (panel a, white lines) should be roughly symmetrical and superimposable, indicating minimal mechanical perturbation due to scanning. (a,b,c): based on the observed signal/noise ratio, an height threshold (green mask in panel a, dashed line in panels b and c) is utilized to separate features subjected to successive analysis steps from the background. A threshold of 10 nm was used in most cases. (d): If present, manifestly non-globular impurities or imaging artifacts are manually excluded from the analysis. Mutually- or edge-touching globular objects are also discarded. For each remaining globule (green mask), the largest inscribable disc is then plotted (white circles), discarding objects having inscribed disc radii below 10 nm. (e): Each remaining object is considered a putative vesicle, and its morphology is parametrized with two quantities measured from its AFM image: the corrected (see Figure S2) radius R_Proj_ of the largest possible disc (white circle) inscribed within the boundary delimited by the height threshold (green dashed line), and the highest Z value occurring within it, H_S_. Scalebar is 75 nm. (f): Geometrical approximation of the spheroid shape of a surface-adhered vesicle with a spherical cap having height H_S_, surface radius A_Cap_ and spherical radius R_Cap_. While H_S_ is always directly measured on the AFM image (see panel e), R_Cap_ and A_Cap_ are calculated from as follows: if R_Proj_ < H_S_, R_Proj_ is taken as a good approximation of R_Cap_; and when R_Proj_ > H_S_, R_Proj_ is taken as a good approximation of A_Cap_. In all cases, contact angle α is then calculated via simple trigonometry calculations (see supporting information).

Putative vesicles are then singled out from the background by marking all pixels exceeding a height threshold (Figure 1a). The employed threshold value was 10nm in all cases except for DOPC samples, for which we employed a 5nm threshold (for reasons explained below). Objects touching any edge of the image were automatically excluded from successive analysis. We then manually excluded objects evidently corresponding to clusters of two or more adjoining globular objects, or to imaging artifacts such as vesicles that detached themselves from the surface between successive scan lines, resulting in non-globular shapes with sharp drops along the slow scan axis (Figure 1d). The radius of the largest possible inscribed disc was then calculated for each object (Figure 1d, white circles); those with an inscribed circle radius <10 nm were discarded to exclude spikes and streaks from successive analysis.

Figure 1e shows a representative AFM image of a single putative DPPC vesicle. Our morphometrical analysis starts with the consideration that shape observed in AFM micrographs is the combination of the vesicle’s true shape, probe convolution, feedback artifacts and the intrinsic AFM limitation of not being able to follow the shape of objects with fractal dimension above 1 along the Z axis [Valle 2017]. Images can be optimized for minimal feedback artifacts (as discussed above), and their quantitative analysis can take probe convolution into account (see Figure S2). The observed AFM morphology is thus assumed to be a close “pseudo-3D” rendition of the examined object, resulting from the combination of the object’s true height values measured along the Z axis and its projection on the XY plane. According to this, a globular object’s true maximum surface height H_S_ and projected surface radius R_Proj_ can be quantitatively measured from its AFM image (Figure 1e): H_S_ is simply its maximum Z value, while R_Proj_ corresponds to its maximum inscribed disc radius corrected for tip convolution (see Figure S2 and S3).

We then assume that the spheroid shape of a surface-adhered vesicle can be approximated to that of a spherical cap [Seifert 1990] with a height equal to H_S_ and a projected surface radius equal to R_Proj_ (Figure 1f). The vesicle’s projected radius R_Proj_ is used as the best approximation of its curvature radius (R_Cap_) if R_Proj_<H_S_ (Figure 1f, left panel); and of its base radius (A_Cap_) if R_Proj_>H_S_ (Figure 1f, right panel). The corresponding vesicle-surface contact angle (α, see Figure 1f) and total membrane area (A_S_, see Figure 2) can be obtained via simple trigonometry calculations (see Supporting Information). Finally, we estimate the vesicle’s size in solution by assuming that even if its shape (originally spherical) was distorted upon interaction with the surface, its membrane underwent negligible stretching [Jackman 2013], thus allowing us to calculate the diameter of a sphere of area A_L_ equal to the A_S_ value recovered from AFM imaging (Figure 2).

**Figure 2.**
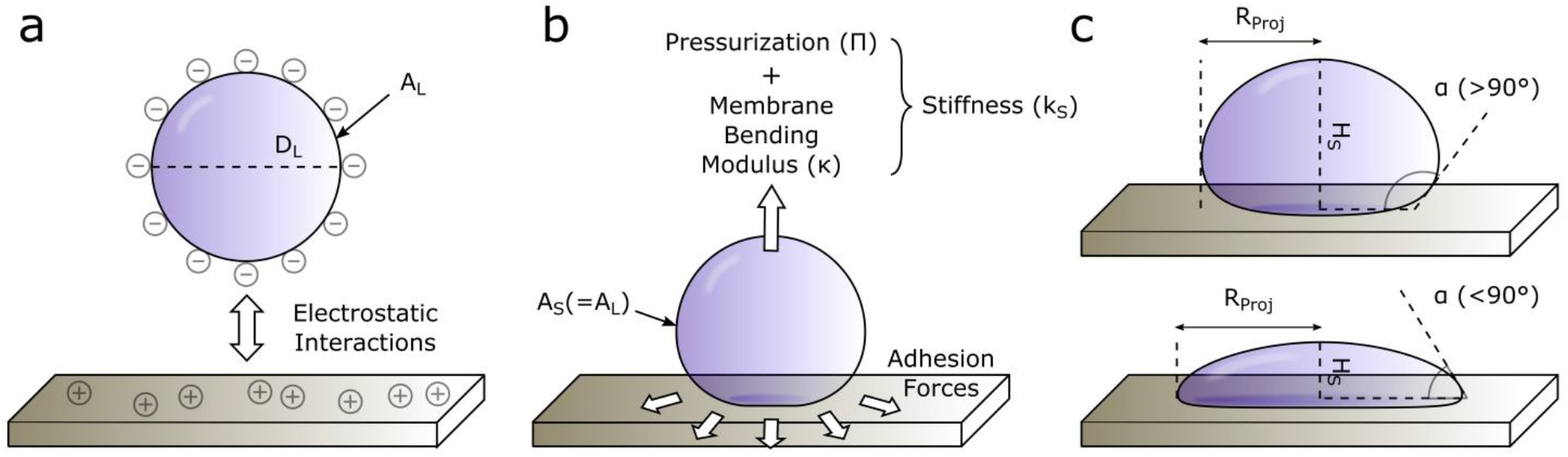
Schematic depiction of the surface adhesion process of a vesicle. (a): In liquid, the vesicle’s average shape is a sphere with diameter D_L_ (‘diameter in liquid’) and corresponding total membrane area A_L_ (‘area in liquid’). All vesicles utilized in this study have a negative ζ-potential in ultrapure water, and are thus electrostatically attracted to substrates coated with poly-L-lysine. (b): When the vesicle first contacts the substrate, adhesive forces tend to maximize surface/membrane contact, causing the deformation of its previously spherical shape into an increasingly oblate spheroid. Membrane stretching is assumed to be negligible throughout the whole process, and thus the total membrane area of the vesicle on the surface (A_S_) is equal to A_L_ (see panel a). The vesicle resists deformation to a degree quantified by its membrane bending modulus κ and internal pressurization Π, which jointly contribute to overall stiffness (k_S_). (c): The equilibrium geometry of the adsorbed vesicle is thus a function of its stiffness k_S_ (see panel b), and can be quantified in terms of height H_S_ and projected radius R_Proj_. These two values can be used to calculate the vesicle’s contact angle α, which describes the entity of its oblate deformation independently from its size; α will be >90° when H_S_>R_Proj_ (top), and <90° in the opposite case (bottom). Comparatively stiffer vesicles will experience smaller deformations and will thus have larger measured α (top) than softer ones (bottom).

### Nanomechanical screening of vesicles via AFM imaging

The rationale for our mechanical screening methodology is schematized in Figure 2. In absence of external perturbations, the average shape of a vesicle in solution is spherical (Figure 2a) and can be geometrically characterized in terms of its diameter (D_L_) and total surface area (A_L_). Most, if not all, EVs have a negative surface charge [Deregibus 2016; Konoschenko 2018; Buzás 2018] and can adhere to positively charged surfaces by electrostatic interactions exerting an attractive force between its membrane and the substrate [Seifert 1990]. Upon interaction, adhesion forces deform the initially spherical vesicle into an increasingly oblate shape. This deformation is opposed by both membrane rigidity and luminal pressurization, which jointly contribute to the vesicle’s observed stiffness (Figure 2b). The extent to which a surface-adhered vesicle is deformed at equilibrium is thus a function of its stiffness, with higher k_S_ values resulting in smaller geometrical distortions and softer vesicles assuming more oblate shapes [Reviakine 2012]. The vesicle-surface contact angle (α) can be employed as a size-independent quantitative descriptor of the adhered vesicle’s deformation (Figure 2c).

With the opportune precautions (see materials and methods), simple AFM imaging in liquid can be used to determine the unperturbed equilibrium geometry of EVs deposited on a substrate in terms of their height H_S_ and surface-projected radius R_Proj_ (Figure 2c). These values can be used to calculate each vesicle’s contact angle α and (assuming membrane area conservation during deformation) its original solution diameter DL.

It is important to note that CH theory assumes κ to be an intrinsic property of vesicles formed by the same type of membrane, while k_S_ is expected to vary with vesicle size [Li 2011]. However, we hypothesize that k_S_ variations observed within populations of vesicles of the same type will be relatively small in the relatively narrow size distribution most relevant to EV research (30-500 nm in diameter). If this is true, populations of compositionally similar vesicles should show a limited dispersion of α values across different vesicle sizes, possibly small enough to resolve their distributions.

### Vesicles of the same type have a characteristic average contact angle value

To verify the above hypothesis on the simplest possible vesicular objects, we first prepared solutions of synthetic liposomes having a negative ζ-potential (DOPC, POPC, DPPC and DSPC) in ultrapure water, deposited them on PLL-coated substrates, captured their adhered morphology with in-liquid AFM imaging, then calculated α and D_L_ values for several hundreds of individual vesicles. For each type of liposome, we plotted the calculated values of all individual vesicles as points on α versus D_L_ graphs (Figure 3). The α values of DOPC and POPC vesicles seem to be weakly negatively correlated with their size, while DPPC and DSPC plots suggest the opposite trend. It is interesting to note that in all cases, most of the deviation from a horizontal, flat distribution occurs in smaller (D_L_<50nm) vesicles, while larger ones seem to converge towards an average α value. Despite these deviations, all the examined liposome types show a relatively narrow global distribution of contact angle values at all observed diameters D_L_, suggesting that the adhesion geometry of a population of vesicles with identical composition can be broadly summarized by their average α value.

**Figure 3.**
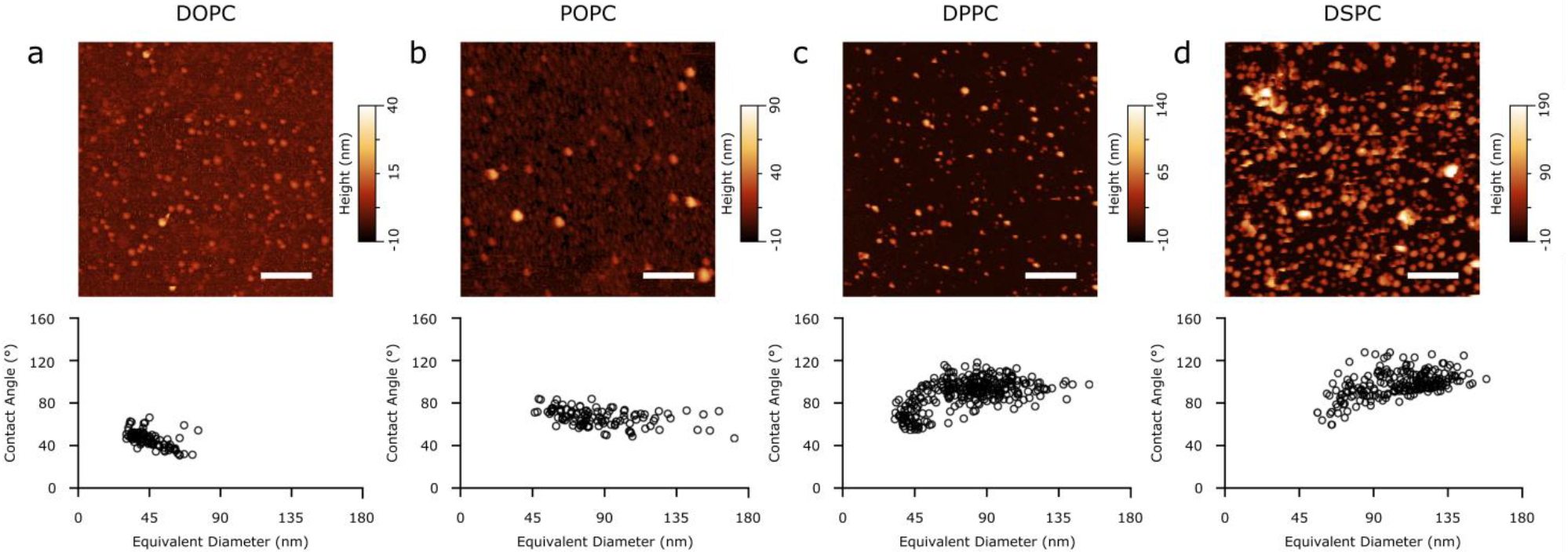
Representative AFM images (top) and Contact Angle vs Equivalent Diameter scatterplots (bottom) of (a) DOPC, (B) POPC, (C) DPPC, and (D) DSPC liposomes. All scalebars are 1μm.

### The contact angle of adhered vesicles is linked to their stiffness

Liposomes in the chosen series (DOPC, POPC, DPPC and DSPC) have increasing κ values [Nagle 2013; Dimova 2014; Yi 2009], which in absence of osmotic imbalances across the membrane result in a correspondingly increasing k_S_ trend. We first verified this assumption via AFM-FS experiments, measuring an increasing trend of κ in the range of 9-20 k_B_T and a correspondingly increasing trend of k_S_ values in the 10-40 mN/m range for the POPC-DPPC-DSPC series, in accordance with previously reported values obtained with this technique [Vorselen 2017; Li 2011]. We then compared these measurements to the image analysis results described above; Figure 4 shows a comparison of the contact angle distributions for each type of liposome. All α values distributions are roughly symmetrical around a median value, which is different for each liposome, and increases along the series. As hypothesized, stiffer vesicles become less oblate than softer ones upon adhesion, and their α values are on average correspondingly larger. All distributions plotted in Figure 4 are significantly different (t-tests, all pairs, P ≤ 0.0001). This suggests that comparing the distribution of contact angles observed via AFM imaging enables the mechanical differentiation of vesicular samples having similar size distributions.

**Figure 4.**
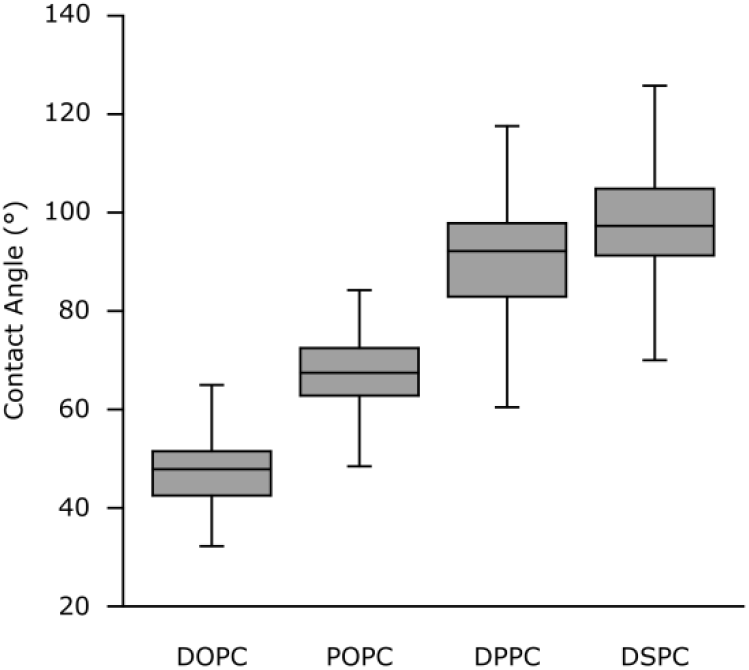
Box plot comparison of the DOPC, POPC, DPPC and DSPC liposomes contact angle distributions. Grey boxes extend between the first (bottom edge) and third (top edge) quartile values, with black lines indicating median values. Whiskers correspond to the lowest (bottom) and highest (top) value found within the distribution. T-tests performed on all pairs of distributions give p-values ≤ 0.0001.

Data reported in Figures 3 and 4 also suggest that the chosen liposome series spans over the entire range of practically measurable α values. DOPC is the softest liposome we could successfully deposit on the employed PLL-functionalized substrates. The size distribution of intact DOPC vesicles on the surface (Figure 3a) is significantly lower than those of the other three liposomes, while it was measured to be similar to that of the POPC sample in solution (see Figure S4), suggesting that larger DOPC vesicles were either ruptured by adhesion forces, or were so compliant as to be mistaken for punctured vesicles and not included in successive analysis. Moreover, even clearly intact vesicles were extremely oblate in shape, with very low H_S_ values. This made it necessary to lower the height threshold used to detect features during image analysis (see materials and methods). The threshold cannot of course be lowered indefinitely, due to intrinsic roughness and instrumental noise; in practice, this sets ~30° as the lowest reliably measurable α values on soft vesicular objects.

At the opposite end of the range, DPPC and DSPC α distributions are substantially overlapping, even if their reported κ values are quite different [Puente 1994; Marsh 2006]. This could be explained by the fact that very stiff vesicles, only experiencing limited deformation upon interaction with the substrate, might have an insufficient contact area to provide stable adhesion; and due to this, they might detach from the surface more readily than softer vesicles when probed by the AFM tip. We indeed observed a high proportion of detachment artifacts (vesicles suddenly ‘disappearing’ in successive scan lines) in DSPC samples. Therefore, the α distribution of DSPC is probably biased toward lower values due to the difficulty of measuring stiffer (and weakly anchored) vesicles.

Taken together, the above considerations seem to imply that negatively charged vesicles having a stiffness between those of DOPC and DSPC should have a practically measurable α range of 30-140° when deposited on PLL-functionalized substrates, and that their average α value should be a function of their k_S_.

### Measuring the contact angle of natural EVs

The same procedure can be applied to samples containing natural vesicles. As reported by Vorselen et al [Vorselen 2018], the mechanical behavior of EVs closely follows that of synthetic liposomes of similar size, even in presence of molecular cargo and integral membrane proteins. Due to this, we expect samples containing a population of EVs with small size and compositional variance to have a correspondingly small α dispersion.

We tested the above hypothesis applying the same procedure used for liposomes on natural EV samples isolated from bovine milk (Figure 5a) and from the parasitic nematode *Ascaris Suum* excretory/secretory products (Figure 5b). Similarly to what obtained with liposomes, the α versus size plots of natural EVs samples (Figure 5a and 5b) show horizontal clusters with no correlation between α and DL, which are indicative of vesicle-like mechanical behavior. This confirms that the purely vesicular nature of the examined samples can be mechanically assessed as previously described for liposomes.

**Figure 5.**
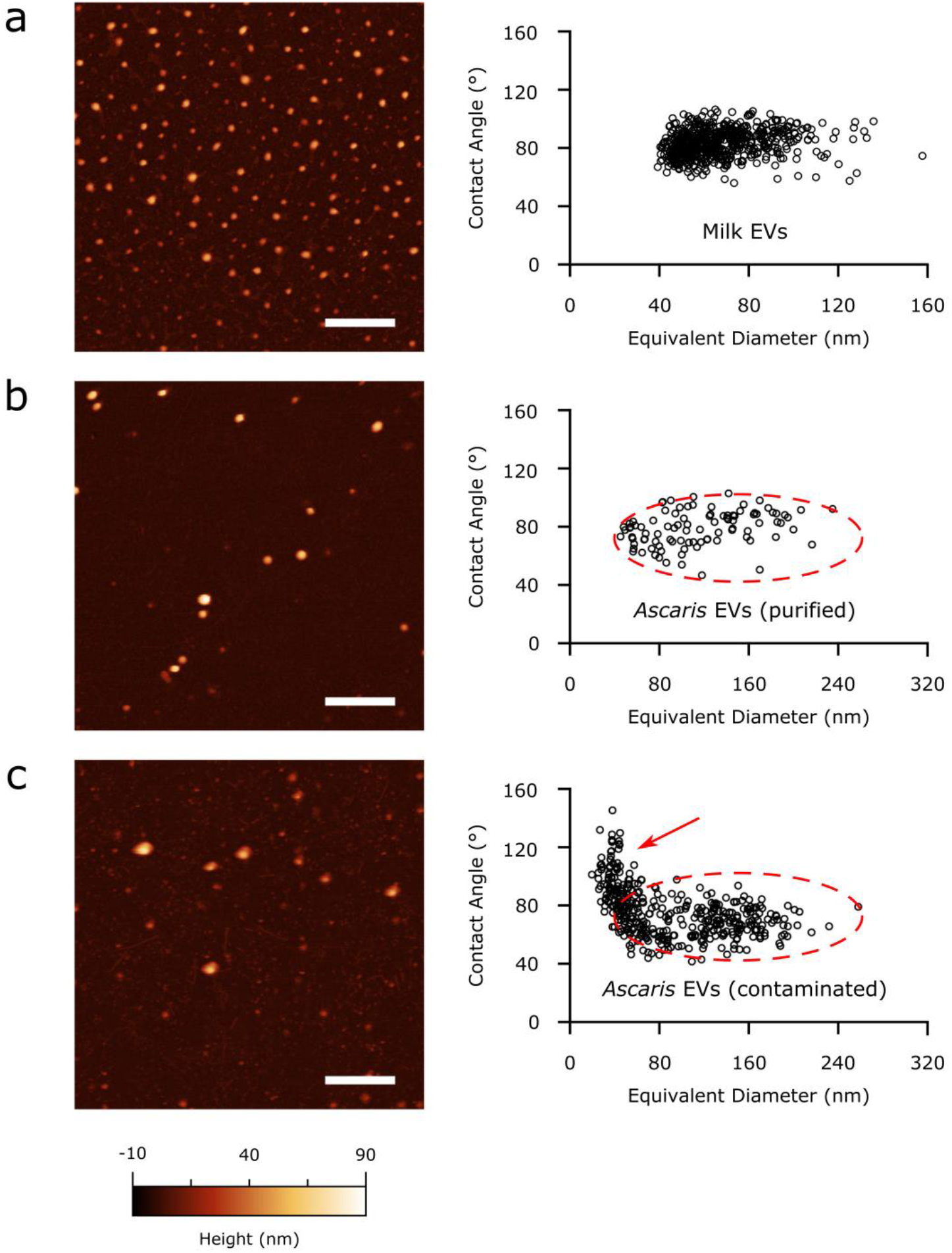
Representative AFM images (left column) and Contact Angle vs Equivalent Diameter scatterplots (right column) of natural EV samples enriched from different sources. (a): EV sample purified from bovine milk. (b): EVs purified from *Ascaris Suum* ES fractions. (c): *Ascaris Suum* EVs contaminated with mycoplasma. All purified EV samples show a relatively small dispersion of contact angles around the same average value at all sizes, resulting in horizontally elongated clusters. Non-vesicular contaminants (red arrow in panel d) do not follow this behavior and appear as additional clusters with large contact angle variations. *Ascaris* EVs in both purified and contaminated samples show in the same zone of the plot (panels b and c, dashed ovals).

Interestingly, the α values of examined natural EVs seem to fall in a relatively narrow range, which corresponds to k_S_ values between those of POPC and DPPC liposomes: 87° ± 7° for bovine milk EVs and 81° ± 10° for *Ascaris* EVs. This observation is compatible with the fact that natural vesicles from different sources can show strikingly similar mechanical properties [Sorkin 2018]. By combining typical EV size constraints (diameter ~40-300 nm) with observed typical EV α values (60°-100°) it is thus possible to draw the boundaries of an area in α vs. D_L_ plots (Figure 6) which could be linked to the presence of “typical” EVs in a sample.

**Figure 6.**
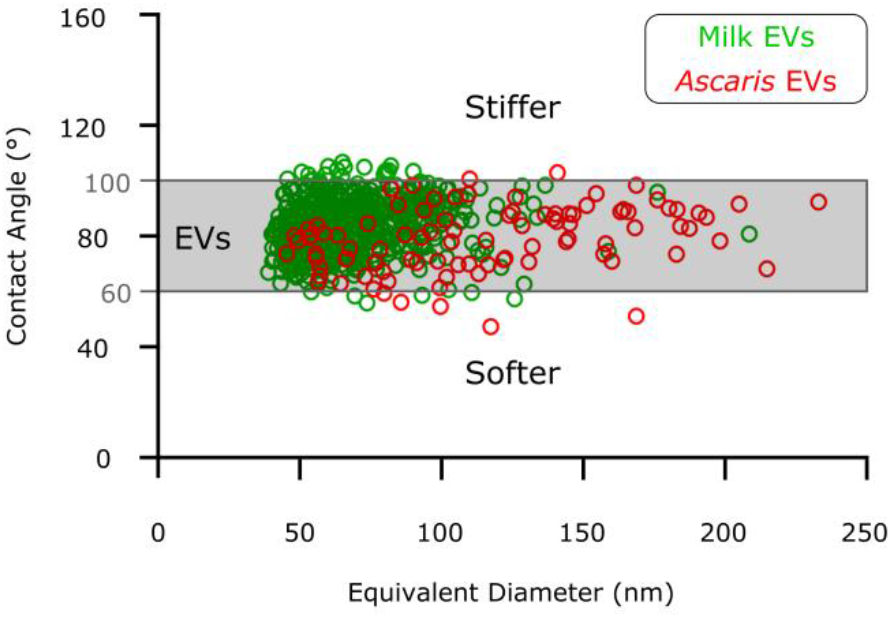
General scheme of a Contact Angle vs Equivalent Diameter plot. The area highlighted in grey delimits values corresponding to ‘typical’ mechanical behavior and size distribution of EVs deposited on a PLL substrate in ultrapure water. Individual EVs from natural sources are plotted together as green (milk EVs) and red (*Ascaris* EVs) circles.

### Contact angle values can be used to discriminate between EVs and impurities

Importantly, EV-enriched samples from natural sources can contain non-vesicular contaminants which could silently bias ensemble-averaged, routine characterization techniques such as *e.g*. dynamic light scattering, ζ-potential, quartz crystal microbalance, flow cytometry, and western blot. Some types of contaminants, having a markedly different morphology from EVs (e.g. membrane patches, fibrils, pili, flagella) can be discerned from EVs by appropriate microscopy techniques, including AFM. However, a purely qualitative visual inspection approach could mistakenly identify as EVs any spurious object having the expected size distribution and a generally spherical shape (e.g. nanosized crystals, protein aggregates, polymer particles). We propose that plotting α versus D_L_ distributions of an EV sample can help in assessing its purity. As discussed above, the α/D_L_ plot of a sample only containing compositionally similar EVs will give a horizontally elongated cluster of points characterized by an average α value. Deviations from this general behavior can be thus taken as indicative of the presence of non-vesicular contaminants.

To test the above hypothesis, we applied our morphometry analysis on a recognized contaminated EV sample. Figure 5c shows the α/size plot of a contaminated *Ascaris suum* EV sample tested with a mycoplasma kit and found positive. The resulting ‘L-shaped’ distribution differs significantly from a corresponding *Ascaris* EV sample tested negative for mycoplasma and bacteria growth (Figure 5b). Besides the expected horizontal band of points with a narrow distribution of α values (which is indicative of vesicles), an additional vertical cluster of objects with a very broad contact angle distribution is present at D_L_ ~ 40 nm. This vertical cluster of points corresponds to globular objects which were included in the morphological analysis because they could not be excluded by qualitative visual inspection alone, but which are mechanically not behaving as a single type of vesicle, thus reflecting non-vesicular contaminants in the sample. The average α value and size distributions of the horizontal clusters of Figures 5b and 5c are comparable (Figure 5b and 5c, red dashed ovals), confirming that the two samples contain the same type of EVs. Our AFM approach can thus distinguish EVs from contaminants on the basis of their mechanical behavior and determine their respective size distributions, facilitating their characterization and successive separation.

### Quantitative estimation of EV stiffness from AFM images

To compare the results of our AFM imaging-based screening with more rigorous, FS-based nanomechanical characterization, we performed AFM-FS experiments (see Figure S1) on a series of increasingly stiffer synthetic liposomes (POPC; POPC:DPPC 1:1 mixture; DPPC; DSPC) deposited on PLL-functionalized substrates, obtaining distributions of their k_S_ values. We then plotted their average α versus average k_S_ (Figure 7), evidencing a strongly linear correlation (R^2^ = 0.97). This suggests that it is possible to quantitatively estimate k_S_ directly from AFM imaging experiments performed on the same substrate used for a calibration line similar to Figure 7.

**Figure 7.**
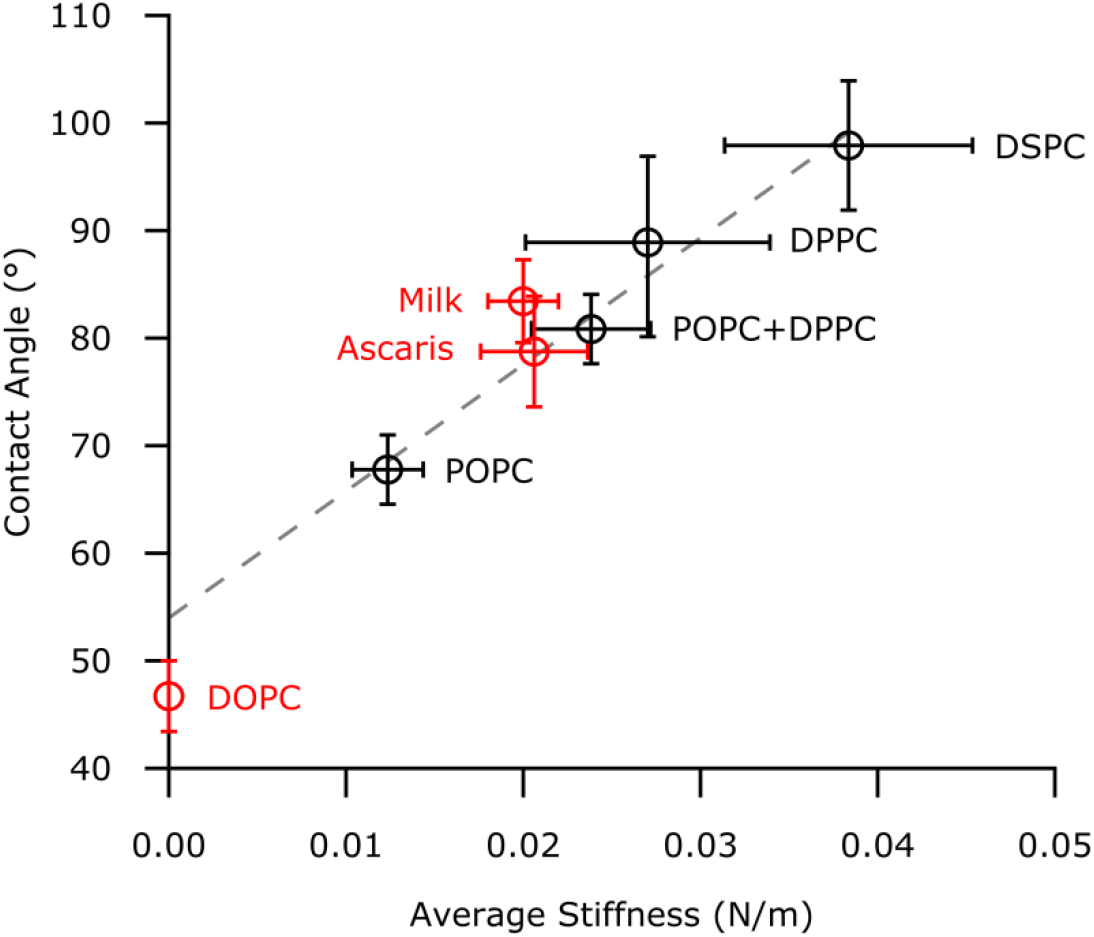
Quantitative correlation between average Contact Angle (α, measured via AFM imaging) and average Stiffness (k_S_, measured via AFM-FS) of vesicles deposited on PLL. Black points correspond to the series of four synthetic liposomes which was used to quantify the α vs k_S_ dependency, showing a strong linear correlation (dashed grey line, R^2^=0.97). Red points correspond to data not included in the linear fit (DOPC and natural EVs). All error bars represent the uncertainties obtained by bootstrapping (1000 repetitions of draws, with replacement). DOPC was plotted at a k_S_ value of zero (see main text). The k_S_ of *Ascaris* and milk EVs (as measured via AFM-FS) is practically coincident with the value obtained by interpolating their average α on the liposome series fit, and in both cases compatible with k_S_ values previously reported for other EVs from natural sources.

It is worthwhile to note that it was impossible for us to perform the full AFM-FS characterization (in terms of k_S_, κ and Π) on some of the samples. In particular, we did not observe measurable linear deformation regimes in any of the DOPC nanoindentation curves, making it impossible to measure its k_S_ via FS. Moreover, we could not measure F_T_ on *Ascaris* and milk EVs, since the vast majority of retraction curves showed complex unfolding/detachment behaviors rather than clean tether elongation plateaus (see Figure S1). Nevertheless, we could easily obtain α/size plots for both samples, and then estimate their expected stiffness values via extrapolation or interpolation of the linear fit shown in Figure 7.

Extrapolating the expected k_S_ of DOPC from its average α yields a nonphysical (negative) value. We interpreted this as a sign of a very low k_S_ value. Interestingly, individual approach/retraction cycles performed on intact DOPC vesicles often show clear tether elongation plateaus on the retraction curve at a specific F_T_, but no linear indentation slope on the corresponding approach curve. This suggests that the low k_S_ of DOPC results in very shallow indentation ‘slopes’ which cannot be distinguished from instrumental noise. Interestingly, if we place DOPC in the α vs. k_S_ plot (Figure 7) by assigning it a k_S_=0, then include it in the linear calibration, the correlation remains highly linear (R^2^=0.98), further reinforcing the observation that α and k_S_ are strongly interdependent across a wide range of values.

We then performed the same α-based k_S_ extrapolation on *Ascaris* EVs, resulting in an expected stiffness value of 21±4 mN/m. In this case however, it was also possible to check extrapolation validity by directly measuring k_S_ via AFM-FS; the experimentally determined stiffness of 20±5 mN/m coincides with the extrapolated value, and is intriguingly similar to previous k_S_ measurements performed on other types of natural vesicles [Vorselen 2018; Li 2011]. *Ascaris* EVs’ experimental point in Figure 7, plotted at their average α (from image analysis) and k_S_ (from FS) is intercepted by the linear fit calculated on synthetic liposomes. The same experimental procedure was then replicated on milk EVs, obtaining strikingly similar results (k_S_ = 20±7 mN/m, see Figure 7). Taken together, these observations suggest that the same strong correlation between α and k_S_ observed in liposomes is also valid for EVs; and that it is thus possible to obtain a quantitative estimate of their stiffness directly from AFM image analysis, without resorting to more time-consuming FS studies. According to this reasoning, the “most typical” natural EV α value of 80° (Figure 6) corresponds to a k_S_ value of ~20 mN/m.

## Conclusion and Perspectives

We have herein described an AFM-based experimental strategy for the nanomechanical and morphological screening of nanosized vesicles. By applying a set of simple experimental precautions and image analysis steps to AFM scans performed in liquid, the proposed procedure makes it possible to discriminate between vesicular and non-vesicular objects in a sample. Furthermore, it allows quantitative size and stiffness estimates for each observed vesicle. Although unable to reach the level of detail afforded by FS-based mechanical assessment methods [Vorselen 2017; Vorselen 2018] previously employed on EVs, the approach proposed here has the advantages of being considerably faster and easier to perform, and of having limited instrumental requirements. Our results also suggest that our approach remains applicable in cases where FS-based approaches might fail.

Being based on the quantitative measurement of contact angles of vesicles adhered to a surface, our method could be extended to other substrates in addition to the PLL-functionalized glass slides employed in this study. This could be functional in modulating surface/vesicle adhesion forces, thus making it possible to better explore vesicles softer than DOPC or stiffer than DSPC by bringing them into the measurable α range or by extending it to the study of positively charged artificial vesicles. Its ability to quickly give a quantitative readout of the interaction between a vesicular object and a nanoengineered surface could be a valid support in developing more quantitative and more reproducible bio-nano material research studies focusing or involving the bio-nano interface [Faria 2018].

While probe convolution has a role in the quantitative assessment of α, its impact is very limited for tip curvature radii below ~15 nm when measuring vesicles within the typical EV size range. However, it could be nonetheless possible to use low quality probes with blunt tips by mixing an internal standard to the sample. This internal standard could be constituted by monodisperse rigid spherical nanoparticles, which could be singled out and used to estimate XY tip convolution, or a synthetic liposome with a previously characterized α value. The latter would appear as an additional horizontal cluster in the α/size scatterplot; apparent R_Proj_ values would then be adjusted by different tip radius values until the reference cluster average α coincided with the expected value.

Our method could in principle also discern between different types of EVs with similar size distributions mixed in the same sample on the basis of their mechanical characteristics. However, EVs examined so far only show minute differences in α and k_S_, thus making their separation only possible by increasing statistical sampling to thousands of individual vesicles, if at all (Figure 6). Of course, we cannot exclude that EVs with putatively more pronounced mechanical differences than those analyzed in our study would be easier to discriminate.

Lastly, the geometrical parameters H_S_ and R_Proj_ can be also used to calculate the volume of each individual adsorbed vesicle in an AFM image. Similarly to how α is linked to k_S_, any measured loss of volume induced by surface adhesion may be linked to Π; it might be thus possible to estimate lumen pressurization without resorting to complex FS experiments. We plan to explore this possibility in forthcoming studies.

## Acknowledgments

This research has received funding from the Horizon 2020 Framework Programme under the grant FETOPEN-801367 evFOUNDRY. PN was supported by a grant from Independent Research Fund Denmark (DFF-6111-00521).

## References

Buzás EI, Tóth EÁ et al. “Molecular interactions at the surface of extracellular vesicles” Semin. Immunopathol. 40, 453–464 (2018).

Calò A, Reguera D et al. “Force measurements on natural membrane nanovesicles reveal a composition-independent, high Young’s modulus” Nanoscale 6, 2275–2285 (2014).

Canham PB “The minimum energy of bending as a possible explanation of the biconcave shape of the human red blood cell” J. Theor. Biol. 26, 61–81 (1970).

Chiang C, Chen C “Toward characterizing extracellular vesiclesat a single-particle level” J. Biomed. Sci. 26, 9 (2019).

Cocucci E, Meldolesi J “Ectosomes and exosomes: Shedding the confusion between extracellular vesicles” Trends Cell Biol. 25, 364–372 (2015).

Deregibus MC, Figliolini F et al. “Charge-based precipitation of extracellular vesicles” Int. J. Mol. Med. 38, 1359–1366 (2016).

Dimova R, “Recent developments in the field of bending rigidity measurements on membranes”, Adv. Colloid Interface Sci. 208, 225–234 (2014)

Faria M, Björnmalm M et al. “Minimum information reporting in bio–nano experimental literature” Nat. Nanotechnol. 13, 777–785 (2018).

Galieva LR, James V et al. “Therapeutic Potential of Extracellular Vesicles for the Treatment of Nerve Disorders” Front. Neurosci. 13, 163 (2019)

Helfrich W “Elastic Properties of Lipid Bilayers: Theory and Possible Experiments” Zeitschrift für Naturforsch. C 28, 693–703 (1973).

Hutter JL, Bechhoefer J“Calibration of atomic-force microscope tips” Rev. Sci. Instrum. 64, 1868–1873 (1993).

Jackman JA, Choi J et al. “Influence of osmotic pressure on adhesion of lipid vesicles to solid supports” Langmuir 29, 11375–11384 (2013).

Jeppesen DK, Hvam ML et al. “Comparative analysis of discrete exosome fractions obtained by differential centrifugation” J. Extracell. Vesicles 3, 25011 (2014).

Konoshenko MY, Lekchnov EA et al. “Isolation of Extracellular Vesicles: General Methodologies and Latest Trends” Biomed Res. Int. 2018, 8545347 (2018).

Krieg M, Fläschner G et al. “Atomic Force Microscopy-based mechanobiology” Nature Reviews 1, 41–57 (2019).

Li S, Eghiaian F et al. “Bending and puncturing the influenza lipid envelope” Biophys. J. 100, 637–645 (2011).

Mardahl M, Borup A et al. “A new level of complexity in parasite-host interaction: The role of extracellular vesicles” Adv. Parasitol. 104, 39–112 (2019).

Marsh D “Elastic curvature constants of lipid monolayers and bilayers” Chem. Phys. Lipids 144, 146–159 (2006).

Montis C, Zendrini A et al. “Size distribution of extracellular vesicles by optical correlation techniques” Colloids Surfaces B Biointerfaces 158, 331–338 (2017).

Nagle JF, “Introductory Lecture: Basic quantities in model biomembranes” Faraday Discuss. 161, 11–29 (2013).

Nečas D, Klapetek P “Gwyddion: an open-source software for SPM data analysis” Cent. Eur. J. Phys. 10, 181–188 (2012).

Ostenfeld MS, Jeppesen DK et al. “Cellular disposal of miR23b by RAB27-dependent exosome release is linked to acquisition of metastatic properties” Cancer Res. 74, 5758–71 (2014).

Paolini L, Zendrini A et al “Biophysical properties of extracellular vesicles in diagnostics” Biomark. Med. 12, 383–391 (2018).

Parisse P, Rago I et al. “Atomic force microscopy analysis of extracellular vesicles” Eur. Biophys. J. 46, 813–820 (2017).

Fernandez-Puente L, Bivas I et al. “Temperature and chain length effects on bending elasticity of phosphatidylcholine bilayers” Europhys. Lett. 28, 181 (1994).

Reviakine I, Gallego M et al. “Adsorbed liposome deformation studied with quartz crystal microbalance” J. Chem. Phys. 136, 084702 (2012).

Roy S, Hochberg FH et al. “Extracellular vesicles: the growth as diagnostics and therapeutics; a survey” J. Extracell. Vesicles 7, 1438720 (2018)

Seifert U, Lipowsky R “Adhesion of vesicles” Phys. Rev. A 42, 4768–4771 (1990).

Shao H, Im H et al “New Technologies for Analysis of Extracellular Vesicles” Chem. Rev. 118, 1917–1950 (2018).

Sharma S, Rasool HI et al. “Structural-mechanical characterization of nanoparticle exosomes in human saliva, using correlative AFM, FESEM, and force spectroscopy” ACS Nano 4, 1921–1926 (2010).

Sorkin R, Huisjes R et al. “Nanomechanics of Extracellular Vesicles Reveals Vesiculation Pathways” Small 14, 1–8 (2018).

Théry C, Witwer KW et al. “Minimal information for studies of extracellular vesicles 2018 (MISEV2018): a position statement of the International Society for Extracellular Vesicles and update of the MISEV2014 guidelines” J. Extracell. Vesicles 7, 1535750 (2018).

Valle F, Brucale M et al. “Nanoscale morphological analysis of soft matter aggregates with fractal dimension ranging from 1 to 3” Micron 100, 60–72 (2017).

Van Deun J, Mestdagh P et al. “EV-TRACK: transparent reporting and centralizing knowledge in extracellular vesicle research” Nat. Methods 14, 228–232 (2017).

van Herwijnen MJC, Driedonks TAP et al. “Abundantly Present miRNAs in Milk-Derived Extracellular Vesicles Are Conserved Between Mammals” Front. Nutr. 5, 81 (2018).

van Niel G, D’Angelo G et al. “Shedding light on the cell biology of extracellular vesicles” Nat. Rev. Mol. Cell. Biol. 19, 213–228 (2018).

Vescovi R, Monti M et al. “Collapse of the Plasmacytoid Dendritic Cell Compartment in Advanced Cutaneous Melanomas by Components of the Tumor Cell Secretome” Cancer Immunol. Res. 7, 12–28 (2019).

Vorselen D, Mackintosh F. et al. “Competition between Bending and Internal Pressure Governs the Mechanics of Fluid Nanovesicles” ACS Nano 11, 2628–2636 (2017).

Vorselen D, van Dommelen SM et al. “The fluid membrane determines mechanics of erythrocyte extracellular vesicles and is softened in hereditary spherocytosis” Nat. Commun. 9, 4960 (2018).

Whitehead B, Wu L et al. “Tumour exosomes display differential mechanical and complement activation properties dependent on malignant state: Implications in endothelial leakiness” J. Extracell. Vesicles 4, 29685 (2015).

Xu R, Rai A et al. “Extracellular vesicles in cancer - implications for future improvements in cancer care” Nat. Rev. Clin. Oncol. 15, 617–638 (2018).

Yáñez-Mó M, Siljander PR et al. “Biological properties of extracellular vesicles and their physiological functions” J. Extracell. Vesicles 4, 27066 (2015).

Yi Z, Nagao M et al. “Bending elasticity of saturated and monounsaturated phospholipid membranes studied by the neutron spin echo technique” J. Phys.: Condens. Matter 21, 155104 (2009).

