## Supplementary materials for "AFM-based High-Throughput Nanomechanical Screening of Single Extracellular Vesicles"

### *Supporting Information: trigonometry calculations*

The contact angle value α is always calculated from $H_S$ and $R_{Cap}$ as follows:

(1) $$\alpha = 90 - \sin^{-1}((R_{Cap} - H_S)/R_{Cap})$$

$H_S$ is directly obtained from AFM images; $R_{Cap}$ and $A_S$ are calculated from $H_S$ and $R_{Proj}$ as follows:

(2) $$\text{If } R_{Proj} > H_S\,,\ R_{Proj} \approx A_{Cap}\ ;\ R_{Cap} = \frac{H_S^2 + R_{Proj}^2}{2H_S}\ ;\ A_S = \pi\left(2R_{Proj}^2 + H_S^2\right)$$

(3) $$\text{If } R_{Proj} < H_S,\ R_{Proj} \approx R_{Cap}\ ;\ A_S = \pi H_S\left(4R_{Proj} - H_S\right)$$

Finally, the vesicle's diameter in solution $D_L$, assuming $A_L = A_S$, is

(4) $$D_L = 2\sqrt{\frac{A_S}{4\pi}}$$

*Supporting Figure S1*

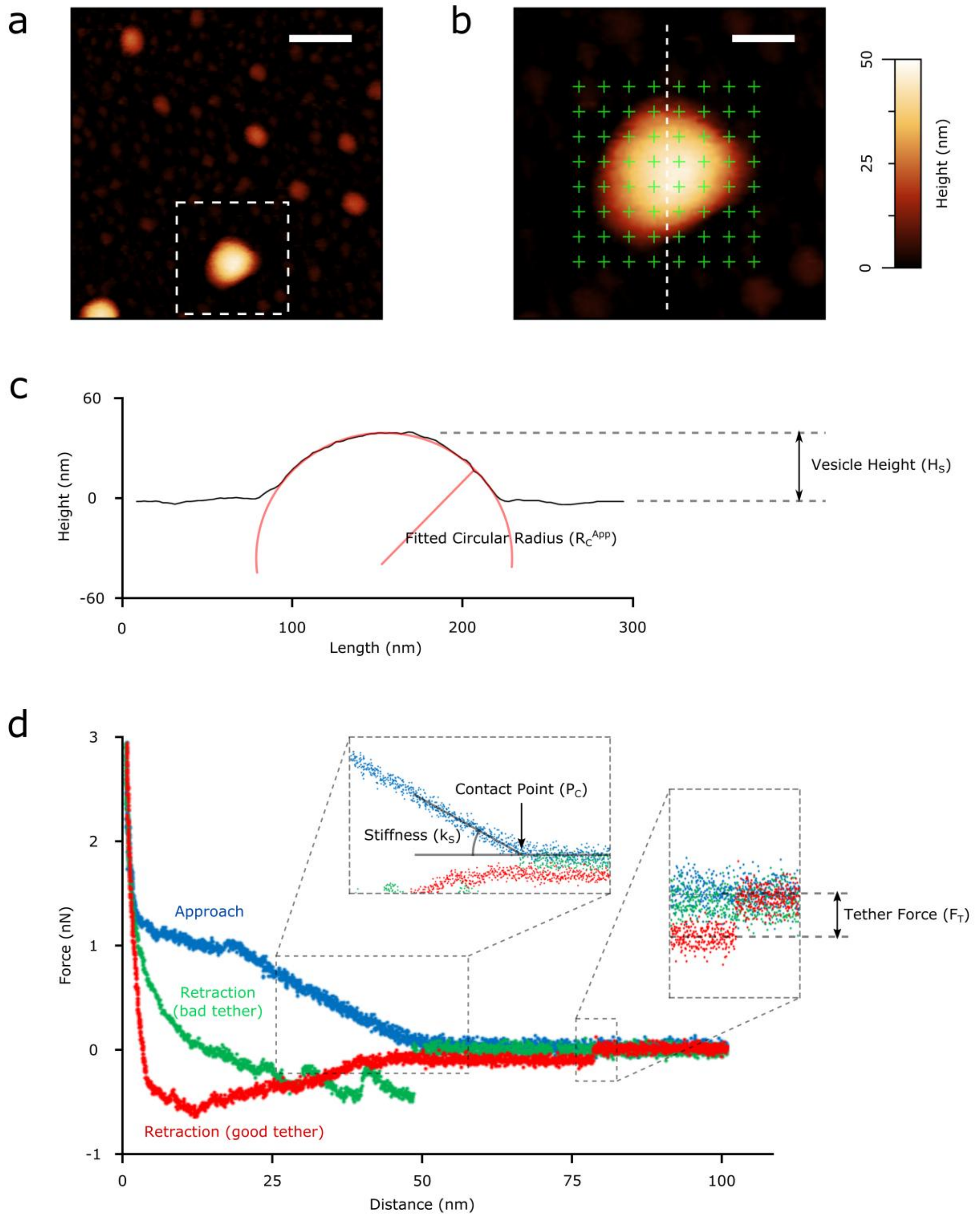


Mechanical characterization of individual vesicles by AFM force spectroscopy was performed following the approach recently suggested by Vorselen et al. [ref Vorselen 2017 in main text]. (a): As a first step, larger areas (scalebar 1µm) were scanned to locate individual isolated vesicles (in this case, a POPC vesicle). (b): The selected vesicle was then imaged at higher resolution (~500x500 nm scan, 512x512 points) to accurately characterize its morphology and to perform force spectroscopy approach/retraction cycles. To avoid intrinsic drifting problems of the piezo and also to gain a more robust estimate of the overall response of the vesicle, multiple indentations were performed following the points on a grid (green crosses) drawn on the vesicle and its surroundings. (c): The height profile along the slow scan axis is fitted with a circle to obtain the curvature radius $R_C^{App}$ which is then corrected for tip convolution (see Figure S2) and used in the normalization of the values of stiffness and internal pressure. Vesicle height $H_S$ is also measured. (d): Typical force/distance curves recorded during approach and retraction FS cycles performed on a single vesicular object. In the approach (blue) curve, applied force remains zero until the tip first touches the vesicle at Contact Point $P_C$, then increases during vesicle indentation. All the curves that showed interactions at Distance values lower than the height $H_S$ observed in the previous imaging step were discarded. According to CHM theory [main text refs Canham 1970; Helfrich 1973], the initial mechanical response of the vesicle to indentation is elastic and linear; the application of a linear fit to this portion of the curve yields the stiffness $k_S$ of the vesicle. The red trace describes the retraction of the tip from the sample and is characterized by the formation of a membrane tether that is pulled by the tip beyond the initial contact point ending with a sharp return to the initial zero force value. The force value measured before this rapid variation is the tether elongation force $F_T$. All the retraction curves that did not resemble the event of tether formation described by the red trace were not considered in the analysis. Obtaining force curves unambiguously showing tether elongations is one of the main issues for the successful application of this FS method to EVs. As exemplified by the green trace, in most retraction traces following the indentation of an EV the presence of abundant membrane proteins and/or peptidoglycans causes the appearance of multiple unfolding/detachment/rupture events (absent in synthetic liposomes) which often avoid the formation and/or identification of single membrane tethers.

### *Supporting Figure S2*

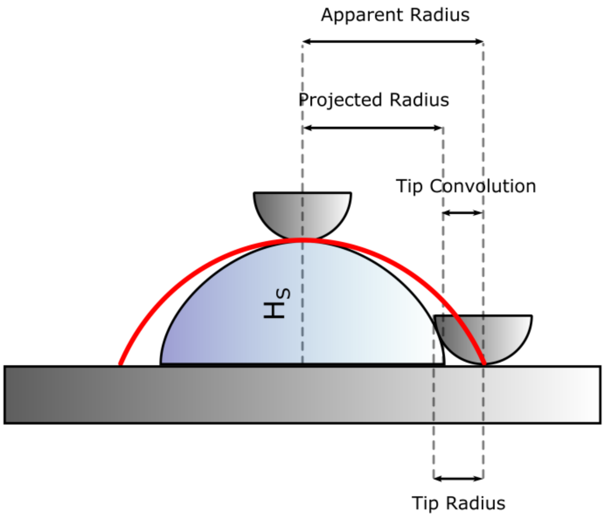


Influence of AFM probe on the nanomechanical characterization of vesicles. The position of a vesicle on a Contact Angle vs. Diameter plot is determined by the $H_S$ and $R_{Proj}$ values measured on its AFM image. While the measurement of $H_S$ is only affected by force setpoint and feedback gain (see main text), $R_{Proj}$ is affected by tip convolution. Due to this, maximum inscribed disc radii values measured on the AFM image should be regarded as 'apparent radii', resulting from the sum of $R_{Proj}$ plus a tip contribution. This tip 'broadening' contribution is variable in entity, and linked to the exact shapes of the tip (its curvature radius at the apex being the most important) and the vesicle (in particular, $H_S$). A clear indication of excessively large tip convolution is a non-circular profile of several vesicular objects in an image; it is not advised to apply image analysis to images where this occurs. Circular profiles (see main text image 2c) can only result from the convolution of two objects having circular shapes along the scanning direction; recording two perpendicular circular profiles on the same object is thus indicative of the fact that the tip is effectively behaving as a hemisphere, and that the largest possible overestimation of $R_{Proj}$ coincides with its apex curvature radius. In any case, the largest broadening effect occurs at the base of the vesicle, which is not included in our image analysis procedure since it only takes into account those portions of objects being above a height threshold (see main text Figure 2a, b and c). This reduces the maximum impact of tip convolution on successive analysis steps to a fraction of the probe's curvature radius. The nominal tip radii of most commercially available 'sharp' AFM tips (e.g. from 2 to 12 nm for the Bruker SNL probes employed in this study) limits the maximum possible overestimation of $R_{Proj}$ to ~10 nm in the worst case scenario for vesicles with $\alpha \geq 90°$. Progressively shallower vesicles would be less affected; the total result being a ~5% underestimation of $\alpha$ for a 'typical' vesicle with $H_S$ = 50 nm in the worst possible scenario.

These considerations suggest that, by using tips with apex curvature radii ≤10 nm and by taking the opportune precautions detailed in the materials and methods section of main text, one can in most cases neglect tip de-convolution. It is important to note that several pieces of information obtained from a Contact Angle vs. Diameter scatterplot (e.g. the presence of contaminants, the attribution of items in a horizontally elongated cluster to vesicle-like mechanical behavior, the relative position of clusters) are unaffected by tip convolution. We only advise tip convolution correction in those cases in which the quantitative readout of $\alpha$ is crucial (e.g. for the quantitative estimation of $k_S$), and if a reliably sharp tip is unavailable. In these cases, it would be possible to correct $R_{Proj}$ values by means of an internal standard, as discussed in the main text (conclusion and perspectives section).

*Supporting Figure S3*

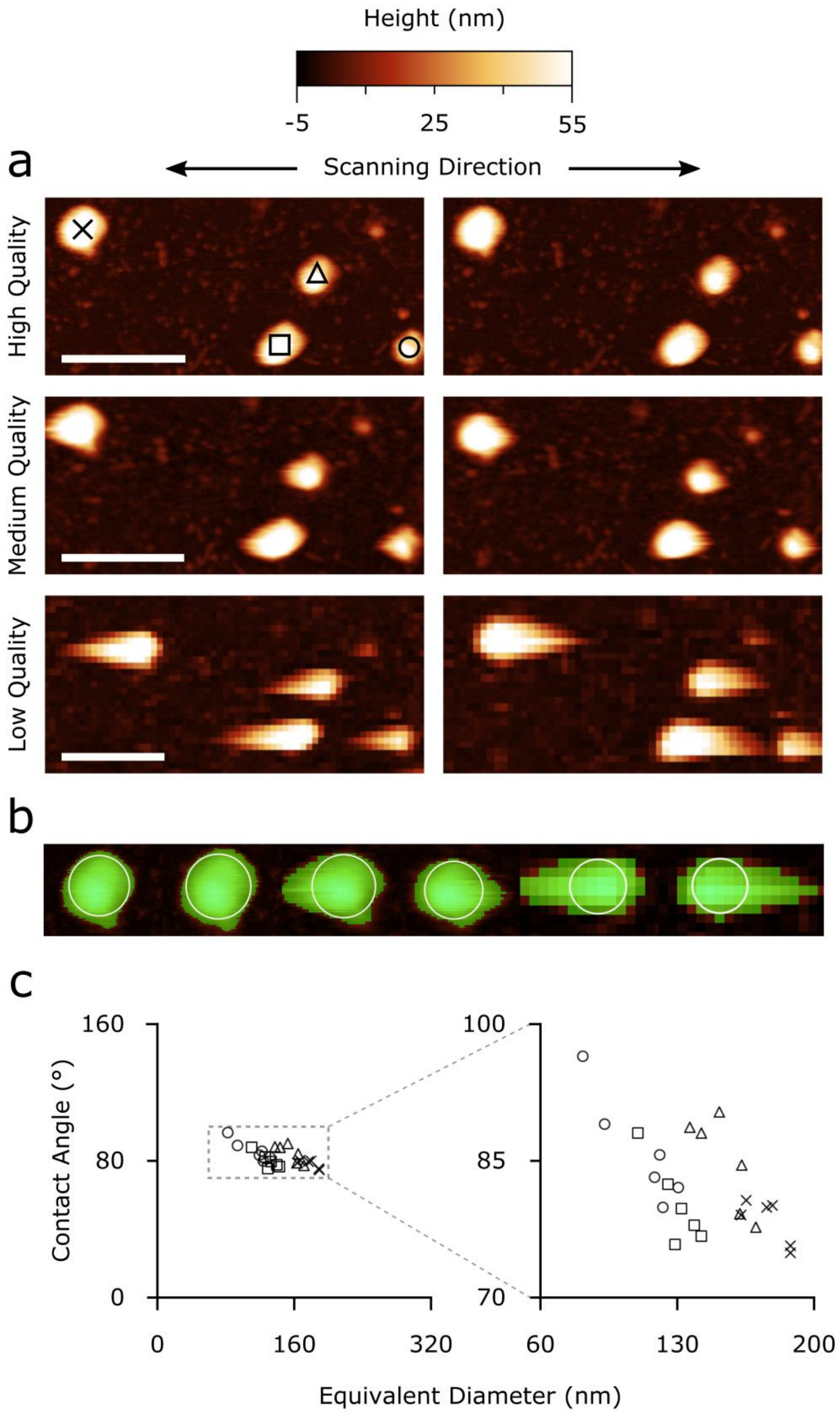


Robustness of the image analysis procedure with respect to imaging quality. Quantitative measurements of α are based on two parameters obtained from AFM images: vesicle height $H_S$ and Projected Radius $R_{Proj}$. While it is of course advised to perform the analysis on AFM images of the best available quality (with respect to resolution, feedback setting, and applied force setpoint: see main text), we found that both $H_S$ and $R_{Proj}$ are only marginally affected by image quality. (a): The two main parameters affecting AFM image quality are feedback gain and resolution (here strictly intended as number of recorded points). We show six

AFM scans on the same four individual *Ascaris* EVs performed in different instrumental conditions. Images in the left and right column respectively correspond to ‘trace’ (toward cantilever apex) and ‘retrace’ (toward cantilever base) fast scan axis directions. Images in the top, middle and bottom rows were acquired with progressively worse imaging quality. Top row: 512x512 points, best feedback setting (vesicles have symmetrical profiles). Mid row: 256x256 points, suboptimal feedback setting (vesicles start to show slow feedback artifacts, elongated ‘tails’ start to appear in the scanning direction). Bottom row: 128x128 points, worse feedback setting (vesicles have long feedback artifacts in the scanning directions). All scalebars are 400nm. (b): Detail of the six scans performed on the vesicle marked with an “X” in panel a. The zone above the selected height threshold is highlighted in green; it is easy to notice how its horizontal deformation caused by feedback artifacts has a limited impact on the maximum inscribed disc radius used to calculate $R_{Proj}$ (white circles). Similarly, reduced resolution has a very limited impact on the maximum height value corresponding to $H_S$. (c): α/size plot of the four vesicles shown in panel a. Each vesicle is marked with the same symbol used in panel a, and plotted at the six slightly different coordinates resulting from image analysis of the six scans of panel a. Pooling the six measurements performed on each vesicle and calculating their variance allows the dispersion of both α and $D_L$ values induced by image quality to be estimated. $R_{Proj}$ and $H_S$ values obtained from the worst images cause deviations of ~5% in α and ~20% in $D_L$ with respect to the best ones.

*Supporting Figure S4*

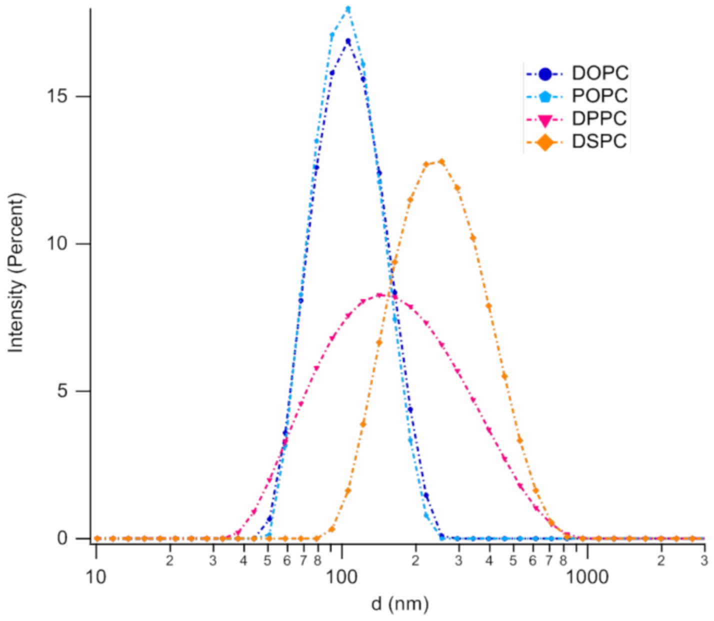


| Lipid composition | Dh (nm) | PDI | Zeta Potential (mV) |
|---|---|---|---|
| **DOPC** | 115 ± 5 | 0.09 ± 0.01 | - 11 ± 3 |
| **POPC** | 110 ± 8 | 0.10 ± 0.05 | - 9 ± 2 |
| **DPPC** | 170 ± 20 | 0.34 ± 0.05 | - 10 ± 3 |
| **DSPC** | 290 ± 20 | 0.20 ± 0.03 | - 11 ± 4 |

Dynamic Light Scattering (DLS) and Zeta Potential characterization of liposomes: (a) Graph reporting the size distribution, as contribution to the scattered intensity, of each lipid composition, as obtained from DLS measurements; (b) Table summarizing average hydrodynamic diameter (Dh) and polydispersity index (PDI) of the lipid vesicles, obtained from DLS analysis, and Zeta Potential values measured for each liposomal dispersion. DLS and Zeta Potential measurements have been performed on a Laser Doppler Micro-electrophoresis (Malvern Zeta Sizer Nano Z), enabling the calculation of electrophoretic mobility and, from this, zeta potential and zeta potential distribution, through the laser interferometric technique M3-PALS (Phase analysis Light Scattering). The measurements have been performed at 25°C. The reported values are an average of three measurements performed on each sample. Measurements were performed at the PSCM Lab (EPN Campus, Grenoble, France). From the reported data it is clearly highlighted that lipid vesicles from DPPC and DSPC (i.e., with a higher stiffness at r.t.) tend to be characterized by larger average sizes and higher polydispersity. All vesicles dispersions in water are characterized by similar, and slightly negative, zeta potential values.

### *EV characterization*

#### *EV preparations purity assessment*

EV preparations form bovine milk and Ascaris suum excretory/secretory products were checked for purity from protein contaminants by the COlorimetric NANoplasmonic (CONAN) assay, which exploits the nanoplasmonic properties of colloidal gold nanoparticles (AuNPs) and their peculiar interaction with proteins and lipid bilayers [Maiolo 2015]. The CONAN assay used in this work consisted of a 6 nM Milli-Q water solution of 14 nm diameter AuNPs. AuNPs were synthesized by the Turkevich's method. The experiments were conducted and data analyzed using the protocols described in [Zendrini 2019]. All the UV/vis/NIR absorption spectra were collected with an Ensight multimode plate reader (PerkinElmer), which allowed collection of the spectra on samples of 100 µL final volume.

The assay consists of an aqueous solution of bare AuNPs at 6 nM concentration. When mixed with pure EV formulations, the AuNPs cluster on the EV membrane, whereas in EV formulations which contain exogenous protein contaminants (EPCs) the AuNPs are preferentially cloaked by such EPCs (an AuNP-EPC corona forms), which prevents AuNPs from clustering to the EV membrane. When AuNPs cluster (are in tight proximity), their localized surface plasmon resonance (LSPR) red shifts and broadens, resulting in a color change of the AuNP solution from red to blue, which can be accurately monitored through UV–vis spectroscopy. The assay red shift is therefore directly related to the purity grade of the added EV formulation and can be conveniently quantified by describing the AuNP UV/vis/NIR absorption spectra with the nanoparticle Aggregation Index (AI), defined as the ratio between the absorbance intensity at the LSPR peak and the intensity at 650+850 nm [Busatto 2018; Mallardi 2018]. For all the analyzed EV formulations separated as described in the main text, the AI values resulted in around 20% of the reference AI of the initial assay (i.e., the dispersed AuNP solution). This proves the EV formulations contained negligible amounts of EPCs. Results reported indicate that the AI for the assayed EV formulation is around 20% of the AI of the starting assay. According to the calibration reported in [Zendrini 2019]. this indicates the samples contain an overall amount of exogenous contaminants < 50 ng/µL.

#### *EV titration*

In the case negligible amount of proteins in EV preparations (< 50 ng/µL) the aggregation index (AI) of the CONAN assay is proportional to the EV number density. We exploited the assay to determine EV total molar concentration (Table ST1) measuring the AI of a POPC liposome calibration curve at known molar concentrations (from 0.8 to 12.5 nM) and interpolating EV AI to the curve. Full details are given in [Busatto 2018 and Zendrini 2019].

### *Supplementary Table ST1*

| EV sample | EV total molar concentration (mol/L) |
|---|---|
| SEC+Myco | 7,07E-08 |
| UC+Myco | 5,64E-08 |
| SEC myco free | 1,00E-08 |
| Bovin milk | 4,25E-07 |

**Table ST1**: EV total molar concentration. UC+Myco: Ascaris suum EV separated from mycoplasma contaminated medium. Separation protocol: Ultracentrifugation (UC). SEC+Myco: Ascaris suum EV separated from mycoplasma contaminated medium. Separation protocol: size exclusion chromatography (SEC). SEC+Myco: Ascaris suum EV separated from mycoplasma free medium. Separation protocol: size exclusion chromatography (SEC). Bovine milk: EV separated from bovine milk.

### *Nanoparticle Tracking Analysis*

EVs size distribution of *Ascaris suum* samples was additionally determined with Nanoparticle Tracking Analysis (NTA). EV separated after ultracentrifugation protocol (UC) or size exclusion chromatography (SEC, fractions 7-10) (see main text for details) were analyzed with a Nanosight NS300 system (Malvern) coupled with a Nanosight syringe pump (Malvern) [using 405 nm wavelength (blue)]. Before any sample analysis, the system was quality checked by measuring a suspension of 100 nm polystyrene beads. In brief, PBS-diluted samples (final volume 1 ml) were injected into the sample chamber using a syringe and the microscope objective was adjusted in order to obtain a clear picture of particles within the beam. Analysis parameter: Flow rate: 10; Temperature: 23°C; Screen gain: 1; Viscosity: Water; Camera level: 10. For each sample, five measurements were performed with a duration of 60 seconds for each repeat/frame. The data were analyzed using NTA software version 3.2.

### *Supporting Figure S5*

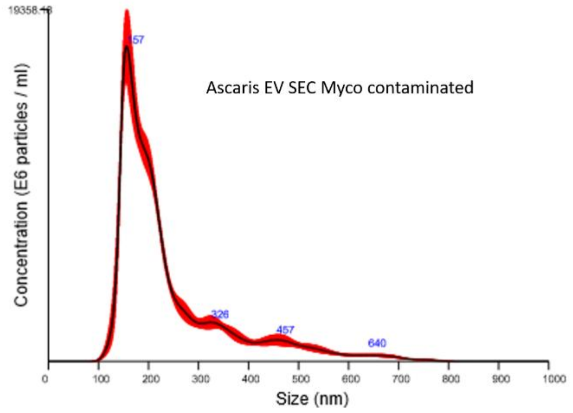


Average finite track length adjustment (FTLA) Concentration/Size graphs for NTA analysis of particles of Ascaris EVs separated with SEC. Medium contaminated with mycoplasma. Red error bars indicate +/- 1 standard error of the mean. Mode 157.5 +/- 3.5 nm

### *Supporting Figure S6*

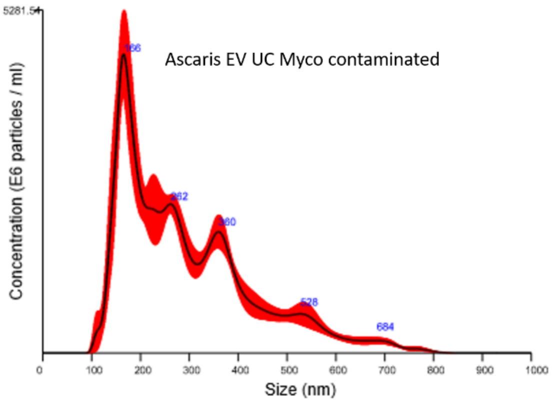


Average finite track length adjustment (FTLA) Concentration/Size graphs for NTA analysis of particles of Ascaris EVs separated with UC. Medium contaminated with mycoplasma. Red error bars indicate +/- 1 standard error of the mean. Mode 164.7 +/- 8.4 nm

### *Supporting Figure S7*

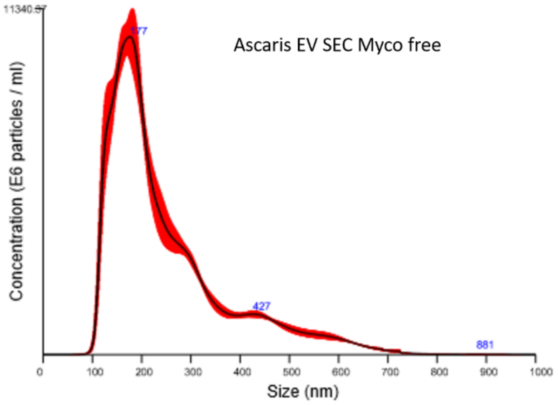


Average finite track length adjustment (FTLA) Concentration/Size graphs for NTA analysis of particles of Ascaris EVs separated with UC. Medium contaminated not with mycoplasma. Red error bars indicate +/- 1 standard error of the mean. Mode 166.4 +/- 6.3 nm

### *EV biochemical characterization*

For biochemical analysis purified EV samples from bovine milk were pelleted at 100000 xg for 65 minutes (in a Beckman Coulter Optima Max-XP with a TLA-55 rotor) in polyallomer microcentrifuge tubes (Beckman) and the pellet was resuspended in sample buffer (62.5 mM  Tris pH 6.8, 2% SDS, 10% Glycerol). Samples were run on a 12.5% SDS-PAGE gel in order to separate proteins. The separated proteins were transferred onto PVDF membranes and blocked in PBS containing 0.2% fish skin gelatin (Sigma-Aldrich) and 0.1% Tween-20. Proteins were detected by immunoblotting using rabbit-anti-human-MFG-E8 (Sigma HPA002807, dilution 1:1000); mouse-anti-bovine-CD63 (BioRad MCA2042G, dilution 1:2000); mouse-anti-human CD9 (Biolegend 312102, clone HI9a, dilution 1:1000); mouse-anti-human-Flotillin (BD 610821, clone 18, dilution 1:500 and the sample was reduced with DTT + β-mercapthoethanol); mouse-a-human-TSG-101

(SC-7964, dilution 1:100 and the sample was reduced with DTT + β-mercapthoethanol); rabbit-anti-bovine Lactoglobulin-β-HRP (Ab112894, dilution 1:1000); rabbit-a-bovine Casein (GTX37769, dilution 1:500). Goat anti-mouse-HRP (Jackson Immuno Research, Suffolk, UK; 1:10000) was used as secondary antibody. HRP conjugated antibodies were detected using SuperSignal West Dura Chemiluminescent Substrate (Thermo Scientific, Landsmeer, Netherlands) and ChemiDoc XRS and Image Lab 5.1 (Bio-Rad) (Figure S8).

### *Supporting Figure S8*

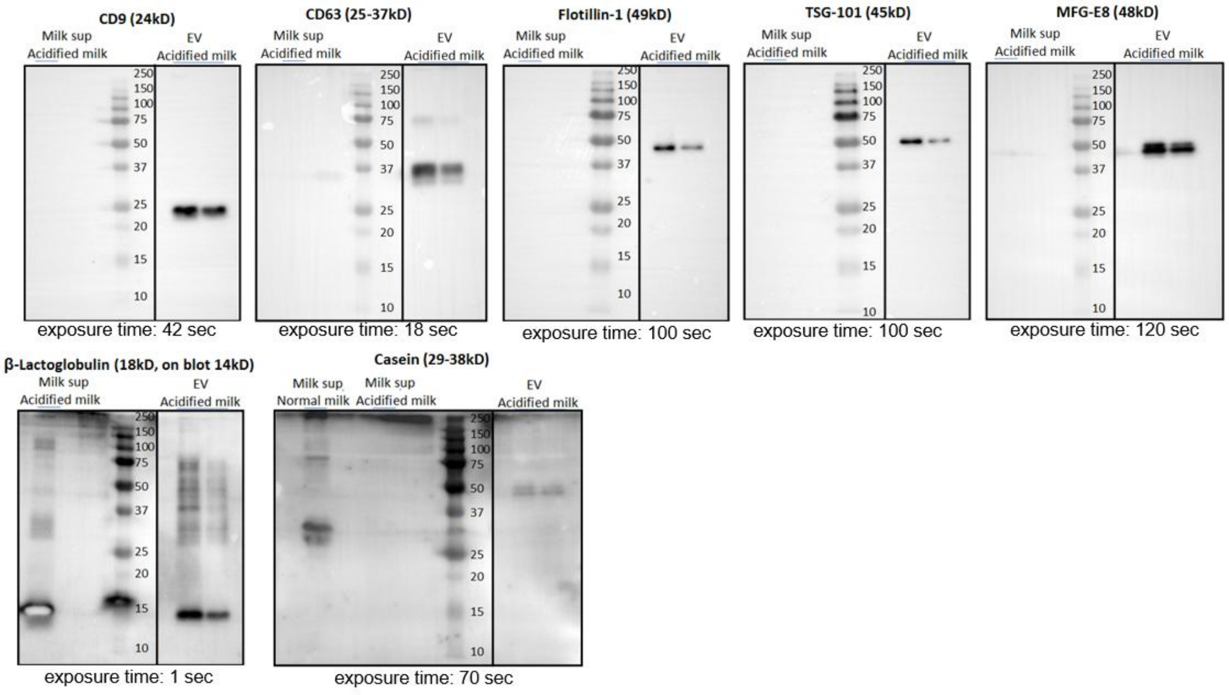


**Figure S8:** Western blot characterization of purified milk EVs for CD9, CD63, Flotillin-1, TSG-101 and MFG-E8. Non EV-enriched proteins β-Lactoglobulin and β casein were included for characterization. For casein, a non-acidified (normal milk) sample was included to show the presence of casein in milk, as compared to acidified milk. Note that β-Lactoglobulin is predicted to be 18 kDa (which was observed in milk supernatant) but the band is lower in EVs. EV samples were technical duplicates as these were isolated from the same raw milk sample.

### *Supplementary References*

[Busatto 2018]: Busatto S, Giacomini A et al. "Uptake Profiles of Human Serum Exosomes by Murine and Human Tumor Cells through Combined Use of Colloidal Nanoplasmonics and Flow Cytofluorimetric Analysis" Anal. Chem 90, 7855-7861 (2018)

[Maiolo 2015]: Maiolo D, Paolini L et al. "Colorimetric nanoplasmonic assay to determine purity and titrate extracellular vesicles" Anal. Chem 87, 4168-76 (2015)

[Mallardi 2018]: Mallardi A, Nuzziello N et al. "Counting of peripheral extracellular vesicles in Multiple Sclerosis patients by an improved nanoplasmonic assay and dynamic light scattering" Colloids Surf B Biointerfaces 168, 134-142 (2018)

[Zendrini 2019]: Zendrini A, Paolini L et al. "Purity and concentration of microliter formulations of EVs by an augmented COlloidal NANoplasmonic (CONAN) method" Front. Bioeng. Biotechnol. (submitted)
